# A bifunctional optical reporter for tracking estrogen response dynamics in neurons

**DOI:** 10.64898/2025.12.14.694224

**Authors:** Alexandra L. Cara, Eartha Mae Guthman, Aarya Mishra, Alexanne L. Hall, Sae Yokoyama, Norma P. Sandoval, Ian Gregg, Krisha Aghi, Kiran K. Soma, Stephanie M. Correa, Annegret L. Falkner, J. Edward van Veen

**Affiliations:** Department of Integrative Biology and Physiology, University of California, Los Angeles, California, USA; Princeton Neuroscience Institute, Princeton University, Princeton, New Jersey, USA; Department of Psychology, University of British Columbia, Vancouver, BC, Canada; Djavad Mowafaghian Centre for Brain Health, University of British Columbia, Vancouver, BC, Canada; Graduate Program in Neuroscience, University of British Columbia, Vancouver, BC, Canada, University of British Columbia, Vancouver, BC, Canada

**Author notes:** Corresponding Authors: J. Edward van Veen, Annegret Falkner, Stephanie Correa. authors contributed equally (Alexandra L. Cara, Eartha Mae Guthman).

**Keywords:** estrogen receptor, viral reporter, estrogen signaling, nuclear receptor

## Abstract

Estrogens are inherently dynamic signaling molecules whose fluctuations induce profound shifts in physiology and behavior. However, methods for tracking the dynamics of estrogen action *in vivo* are limited and not tissue-specific, making it difficult to directly relate dynamics to ongoing physiological, behavioral, and neural changes. Traditionally, estrogen sensitive cells have been identified by metrics such as receptor expression or radioactive ligand binding, but these measures do not capture the dynamic nature of hormone response. In addition, current methods for assessing the dynamics of estrogens in an individual, including microdialysis and vaginal cytology, lack temporal and spatial resolution.To address this need we developed a bifunctional reporter called neuro-seeER (neuronal see Estrogen Response) that can be used to track changes in exogenous and endogenous estrogen dynamics *in vitro* and *in vivo,* including longitudinally in behaving animals. We confirmed that the fluorescent response requires estrogen receptor expression, is specific to activation with estradiol, and is consistent with the timescale of transcriptional induction. Neuro-seeER expression in the hypothalamus of Esr1-Cre mice revealed dynamic labeling of estrogen-responsive cells following treatment with exogenous estradiol, and fluorescence-aided cell sorting followed by RNA-sequencing confirmed that high neuro-seeER response is associated with enriched expression of genes associated with endogenous estrogen responses. Using a novel “snapshot” photometry method to track the dynamics longitudinally *in vivo,* we demonstrate that we can robustly detect exogenous estrogen response across multiple hypothalamic sites simultaneously. Finally, we demonstrate that this tool is uniquely suited to capturing endogenous estrogen response dynamics *in vivo* across the long timescale of the estrous cycle, revealing individual differences in neural estrogen dynamics. Together, these findings reveal unexpected cellular and temporal heterogeneity of the transcriptional response and demonstrate the feasibility of tracking the dynamics of hormonal response alongside additional neural, cell signaling, or behavioral measures.

## Main

Estrogens are highly dynamic hormones, and their changing levels drive striking transformations in both physiology and behavior^1–9^. It is currently difficult to track their effects continuously *in vivo,* hampering progress in understanding the dynamic role that estrogens play in updating neural activity and cell state. Estrogen action is typically inferred through measurement of hormone levels in circulation and detection of receptors in target tissues, but these methods ignore critical variation in site-specific synthesis, time scales of receptor signaling, and heterogeneous responses across tissue and cell types^10–13^. In addition, circulating estrogen levels are a poor proxy for the *effects* of estrogen, since multiple factors, including the dynamic expression and post-translational modification of nuclear receptors and co-factors, may influence cellular signaling (reviewed in^14^). Moreover, measures of estrogens in circulation are not well-suited technically for integration with *in vitro* or *in vivo* measurements of neural activity. Therefore, there is an urgent and unmet need to develop novel optical tools that can be used to dynamically track estrogen signaling in various brain sites and across timescales from minutes to months.

One approach to detecting a functional estrogen response is to assess mechanisms downstream of the receptor. These can be evaluated via molecular metrics such as kinase cascade phosphorylation and binding of estrogen receptors to estrogen response elements (EREs) or by cell biological effects such as cell proliferation or actin remodeling. Type I nuclear hormone receptors, including estrogen (ERα, ERβ), androgen (AR), and progesterone receptors (PR), undergo conformational changes when bound to their respective ligands, inducing dimerization, and binding to hormone response elements (HREs) in the promoter or enhancer regions of target genes, ultimately modifying transcription^15^. Previous approaches for studying estrogen response have demonstrated that EREs placed upstream of a minimal (inducible) promoter can activate reporter expression in response to estrogens. These strategies have elegantly shown varying estrogen activity across different organs and begun to explore the dynamics of estrogen response. The most widely used reporter mouse (ERE-Luc^16,17^) relies on luciferase-based assays, which, although sensitive, are limited by their need for a substrate and cofactors. A similar ERE-βgal mouse (ER action INdicator (ERIN)^18^) has been used to examine the effects of selective estrogen receptor modulators, but is also limited by the same need for substrate, as well as an inability to report estrogen activity in living tissues. Another similar ERE-EGFP mouse^19^ does not require substrate, but uses a non-canonical ERE sequence and a fluorophore with a 24h half-life, both complicating the analysis of estrogen response dynamics.

Due to the limitations of previous tools, the dynamics of estrogen response in the estrogen sensitive nuclei of the hypothalamus have not been examined longitudinally in behaving animals. In addition, previously developed tools lack the ability to resolve estrogen response with cell-type specificity. Given recently appreciated heterogeneity in gene expression and function of estrogen responsive neurons^20–25^, we propose that examining estrogen signaling within restricted populations of neurons will be critical to understanding its pleiotropic effects on physiology and behavior. Furthermore, the available tools do not provide a way to identify neurons capable of response, defined by expression of estrogen receptors (estrogen sensitivity), but not actively responding (estrogen responsivity).

To overcome limitations of previous methods, we developed an improved dual function reporter, termed a seeER (“see Estrogen Response”). The seeER strategy labels Cre-expressing cells with the red fluorophore, FusionRed, and estrogen responsive cells with a destabilized green fluorophore with a 2 h half-life (d2EGFP^26^). FusionRed is driven by the human synapsin promoter, giving neuron specific expression of the Cre-dependent fluorophore. In Esr1-Cre mice, the neuro-seeER fluorescently labels both estrogen sensitive and estrogen responsive neurons, further refining cell-type identification and classification of estrogen responsiveness. Here, we demonstrate how this reporter can be used *in vitro* and *in vivo* to track responses to exogenous and endogenous estrogen fluctuations. We demonstrate how this can be used with standard “off the shelf” photometry, sequencing, and microscopy techniques, making it ideally suited for integration with a wide variety of experimental setups. Further applications using the seeER may involve single cell and/or spatial transcriptomics, combined fiber photometry recording of real time estrogen response and neural activity using genetically encoded calcium indicators, or the examination of estrogen response in molecularly defined neurons across the brain and body.

## Results

### See-ER expression *in vitro* is ligand-specific, receptor-specific, and dynamic

We engineered the seeER to simultaneously report estrogen response with low-latency and also report neuronal cell type information with temporal stability. To achieve these two functions, we made a bicistronic adenoviral vector harboring a Cre-dependent hSyn-Flex-FusionRed cassette and an 8xERE-minimalCMV-d2EGFP cassette (pAAV-hSyn-FLEX-FusionRed-mCMV-8xERE-d2E GFP, Fig. 1A, Fig. S1A). The seeER strategy labels Cre-expressing cells with the red fluorophore, FusionRed, and estrogen responsive cells with a destabilized green fluorophore with a 2 h half-life (d2EGFP^26^). To validate the ligand and receptor dependent expression of EGFP, we tested a nearly identical seeER reporter (pAAV-CMV-FLEX-FusionRed-mCMV-8xERE-d2EGFP, Fig. S1B) in human 293T cells. Using receptor co-transfection, steroid ligand treatment, and multi-well based time-lapse microscopy (Fig. 1B), we imaged and quantified the dynamics of ERE-dependent EGFP expression *in vitro.* As expected, treatment with 17β-estradiol (E2) induces a transient increase in the number of EGFP-expressing cells when the seeER is co-transfected with constructs that confer expression of ERα (Fig. 1C-F) or ERβ (Fig. 1G) but not with constructs that confer expression of the glucocorticoid receptor (GR, Fig. 1C-F) or androgen receptor (AR, Fig. 1H, I). In cells co-expressing seeER and ERα, EGFP expression was induced by E2 treatment (Fig. 1C-F) but not by 5α-dihydrotestosterone (DHT) treatment (Fig. 1H, I). Interestingly, a small number of EGFP+ cells were induced by DHT treatment (Fig. 1H, I) but not E2 treatment (Fig. 1G) in AR-expressing cells, suggesting a low level of ERE-dependent transcription induced by ligand-bound androgen receptors. Together, these findings indicate that the combination of estrogen receptor and an estrogen is both necessary and sufficient for the robust expression of EGFP from the seeER.

**Fig. 1:**
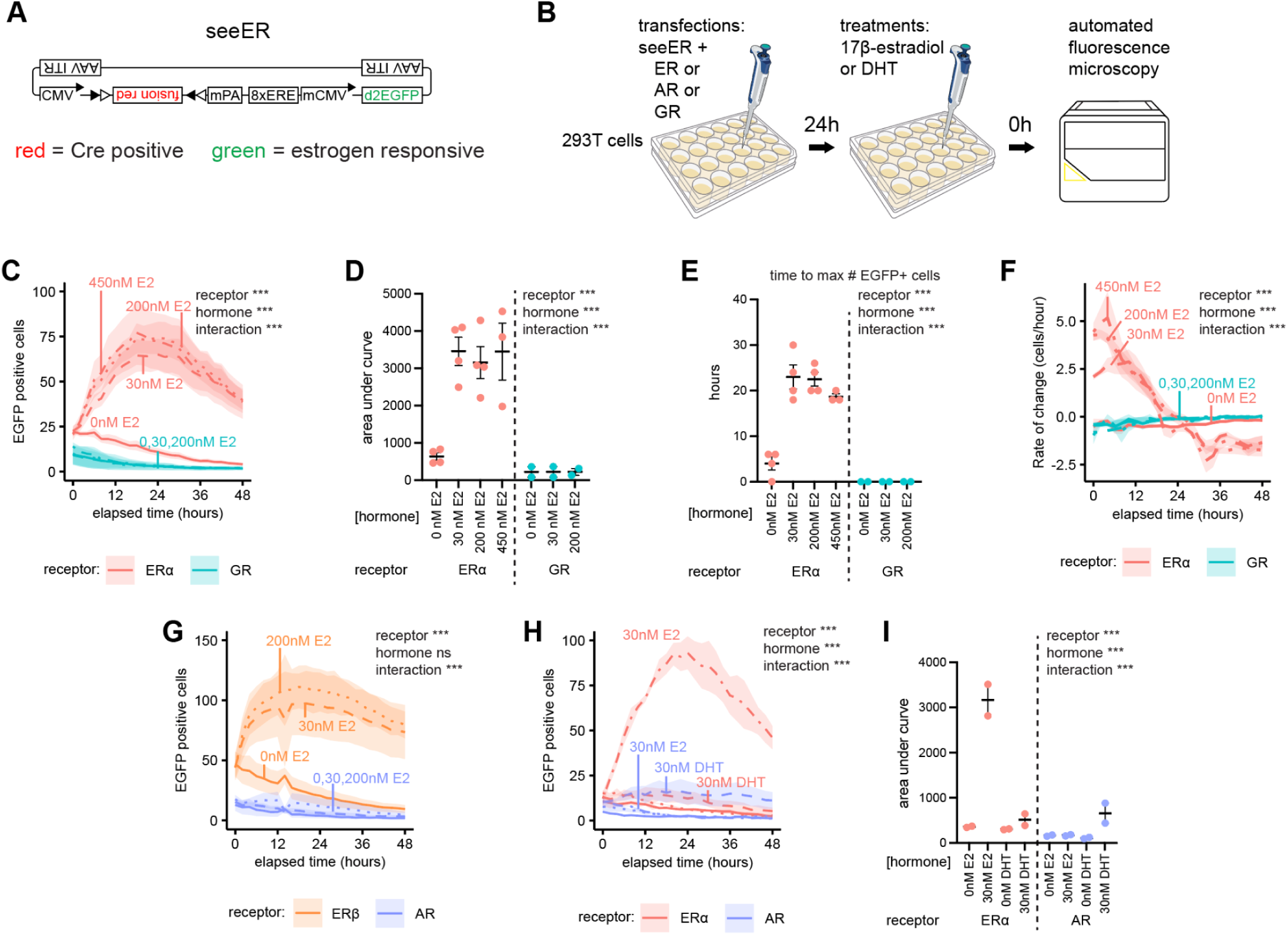
seeER validation *in vitro*. A, seeER reporter that drives Cre-dependent red fluorescence and estrogen-dependent green fluorescence. Eight tandem estrogen response elements (8xERE) act as a binding site for ligand-estrogen receptor (ER) complexes, which drive transcription of a destabilized (2 h half-life) enhanced green fluorescent protein (EGFP). B, workflow for testing ligand and receptor dependence of seeER based green fluorescence using automated fluorescence microscopy (Incucyte). C, Induction of EGFP expression in 293T cells transfected with steroid receptors. Line color indicates receptor transfected, line style indicates hormone treatment. EGFP positivity determined by Incucyte instrument. D, Area under curve analysis spanning 48 h of data shown in panel C. E, Kinetics of appearance of EGFP in response to E2 addition in 293T cells transfected with steroid receptors. F, First derivative analysis of the rate of change of estrogen responding cells. G-H, Induction of EGFP expression in 293T cells transfected with steroid receptors. Line color indicates receptor transfected, line style indicates hormone treatment. I, Area under curve analysis spanning 48 h of data shown in panel H. For all panels, lines and surrounding shading represent mean +/- standard error. Individual points represent biological replicates. Biological replicates were taken as repetitions of the same experiment in different stocks of cells with different passage numbers. See Table S1 for N’s, ANOVA details, and expanded pairwise comparisons.

We compared the effects of E2 on ERα expressing cells and observed that EGFP expression did not differ across the tested dosages. The number of EGFP+ cells was similarly induced by 30 nM, 200 nM, and 450 nM E2 treatment, compared to vehicle (Fig. 1C, D), suggesting that the number of estrogen-responsive cells is not altered beyond a permissive dose. Similarly, the time to the maximum number of EGFP+ cells (Fig. 1E) and the rate of change in the number of EGFP+ cells (Fig. 1F) were similar following 30 nM, 200 nM, and 450 nM E2 treatment, suggesting that the temporal dynamics of the seeER are similar across the range of doses tested.

### Neuro-seeER expression in the hypothalamus is dynamically induced by estrogen treatment *in vivo*

To test whether seeER could be used *in vivo* to label and identify estrogen responsive neurons using standard microscopy techniques, we quantified the seeER fluorescence at specific timepoints following an exogenous estrogen treatment. AAV encoding the neuro-seeER was injected into the medial preoptic nucleus (MPN) and surrounding medial preoptic area (MPO) of ovariectomized (OVX) Esr1*-*Cre mice (Fig. 2A). In female rodents, the MPN and surrounding MPO abundantly express ERα^27^ (Fig. 2B). Following recovery from OVX, mice were injected with either estradiol benzoate (EB; 1 μg, s.c.) or a vehicle control (oil, s.c.). Confirming the effectiveness of the EB treatment, uterine mass, a common bioassay of relative estrogen levels, was higher at 4h, 6 h, and 16h following EB injection (Fig. 2C). When comparing MPO sections taken from vehicle injected mice (Fig. 2D) to EB injected mice (Fig. 2E), the majority of FusionRed+ neurons appear largely similar. However, in EB injected animals a population of bright green neurons appeared (Fig. 2Ei, ii) indicating that a subset of ERα-expressing neurons has responded to the injection of EB. To quantify this effect, EGFP fluorescence was standardized to FusionRed fluorescence as an internal control. EGFP signal relative to FusionRed signal was determined for each individual cell using residuals from a linear regression of FusionRed pixel intensity (independent variable) and EGFP pixel intensity (dependent variable). To account for variability across experiments and tissue sections, the residuals were standardized using z-scores. A standardized residual of 8 was chosen as the cutoff for the definition of “bright” relative EGFP fluorescence (henceforth “estrogen responsive neurons”). This stringent cutoff was used for all microscopy quantification, but standardized residuals of 3-11 reveal significant effects (*p* < 0.05 vs control, Table S2) of EB injection on the number of estrogen responsive neurons. As observed *in vitro*, estrogen treatment led to a significant increase in the number of estrogen responsive neurons at 6 and 16 h after treatment with EB, compared to treatment with vehicle (Fig. 2F). As a control, we analyzed standardized residuals to identify neurons in which FusionRed fluorescence was relatively brighter than EGFP and did not identify patterns of induction or downregulation with EB treatment, confirming that EB treatment selectively altered EGFP expression (Table S2).

**Fig. 2:**
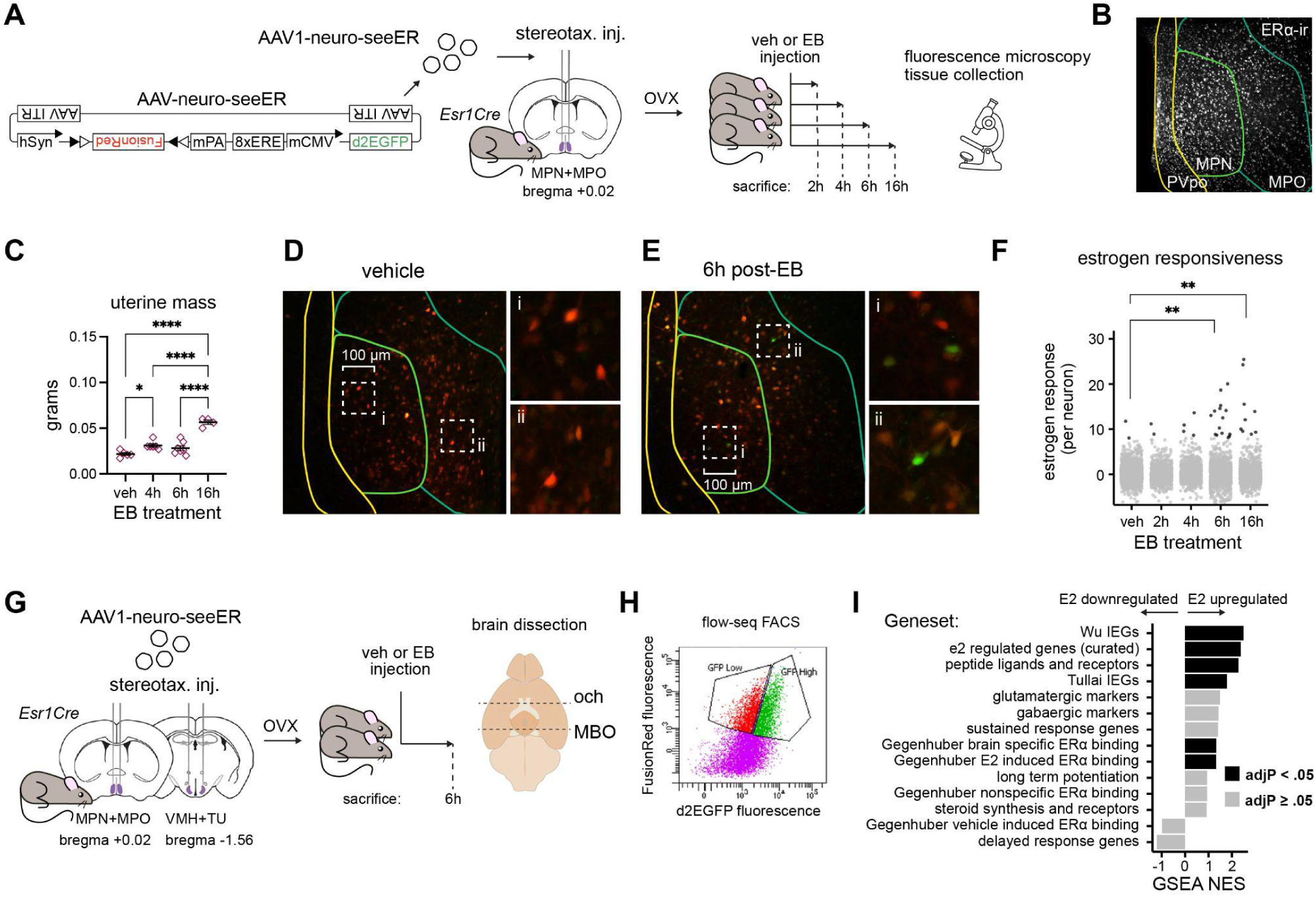
neuro-seeER validation by terminal sampling fluorescence microscopy and flow-seq. A, Bilateral stereotaxic injection of AAV1 harboring neuro-seeER reporter. Mice were sacrificed 2h, 4h, 6h, or 16h after vehicle or EB treatment. B, ERα expression within hypothalamic nuclei at Bregma +0.02 mm, including the regions of interest for quantification. C, Uterine masses of mice used in panels D-F. D-E, ERE-dependent EGFP expression (green) and FusionRed (red) in the MPN+MPO of vehicle (D) or EB (E) treated mice after 6 h. Neurons that display relatively high green fluorescence are shown in insets (i,ii). F, Magnitude of estrogen response in non-responsive (gray) and responsive (green) neurons in the MPN+MPO of vehicle and EB injected animals at indicated timepoints. G, Experimental design to isolate estrogen sensitive neurons from single cell suspension pooled from the MPN+MPO and VMH+TU of adult female Esr1-Cre mice for transcriptomic analysis. Brains were blocked ventrally and trimmed on the coronal plane rostral to the optic chiasm (och) and rostral to the mammillary bodies (MBO). H, Representative flow plot shows sorted cells that were selected for red fluorescence, and then segregated for relatively higher (GFP high) or lower (GFP low) expression of EGFP. I, Gene set enrichment analyses of pathways tested against differentially expressed genes in low vs. high EGFP expression. Positive normalized enrichment scores (NES) indicate pathways coordinately upregulated in high EGFP neurons compared to low EGFP neurons. Black bars indicate that gene set enrichment results are significant with adjusted p-value < .05. See Table S1 for N’s, ANOVA details, and expanded pairwise comparisons.

To test whether SeeER could be used to profile estrogen responsive neurons, we analyzed estrogen responsive neurons using flow cytometry followed by bulk RNA sequencing (flow-seq). In a separate cohort of OVX Esr1-Cre female mice, neuro-seeER was introduced to the MPN+MPO and ventromedial and tuberal hypothalamic nuclei (VMH+TU) by stereotactic microinjection. Regions expressing FusionRed were collected 6 h after treatment with either vehicle (oil, s.c) or EB (1 μg, s.c) (Fig. 2G). FusionRed+ neurons were sorted into estrogen responding and non-responding populations using a linear regression of EGFP and FusionRed fluorescence (Fig. 2H). Injection of 1ug EB did not result in an observable shift in green fluorescence by FACS compared to vehicle and so treatment groups were pooled. Comparing the transcriptomes of estrogen responding vs non-responding neurons confirmed several expected changes in gene expression (see Table S3 for all differentially expressed genes). Estrogen responding neurons were enriched for known estrogen receptor targets, including *Pgr* (LogFC = 1.6, adjusted p = 0.05), *Igf1* (LogFC = 2.26, adjusted p = 0.00049), and *Oxtr* (LogFC = 1.8, adjusted p = 0.002). Gene Set Enrichment Analysis (GSEA, see Table S3 for all gene sets tested) confirmed that estrogen responding neurons have higher overall expression of known hypothalamic estrogen targets (Fig. 2I, Fig. S2A, NES = 2.38, adjusted p = 9.26e-9). GSEA also confirmed that estrogen responding neurons show upregulation of genes that were previously identified as estradiol dependent targets of ERα in the brain (Fig. 2I, NES 1.32, adjusted p = 0.002)^28^. GSEA also found that immediate early genes from two independent studies^29,30^ were significantly enriched in estrogen responding neurons (Wu et al. NES: 2.5, adjusted p value 6.2e-24, Fig. 2I, Fig. S2B), consistent with the established effects of estradiol on hypothalamic neuronal activity^31,32^. Interestingly, the differences in estrogen responses were not associated with significant differences in the expression of genes that encode ERα (LogFC = 1.17, adjusted p = 0.22) or ERβ (LogFC = -0.13, adjusted p = 0.96), confirming that estrogen response cannot be captured by examining receptor expression alone. In addition to expected differences, there were unexpected changes in gene expression including in two UDP-glucuronosyltransferases (Ugt8a LogFC = -1.9, adjusted p-val = 9.1e-18; Ugt1a7 LogFC = 1.5, adjusted p-val = 0.047), genes whose products have been implicated in neurosteroid synthesis and the modulation of estrogen levels in the brain^33^. Estrogen responsive neurons also show surprising changes in expression of key stress hormone genes including the glucocorticoid receptor (*Nr3c1*, LogFC = -0.52, adjusted p = 0.012), and corticotropin releasing hormone binding protein (*Crhbp*, LogFC = 4.32, adjusted p = 6.74e-19). Flow-seq analysis of estrogen responsive neurons from the larger pool of ERα expressing neurons thus identified a broad range of estrogen signatures that were previously identified by multiple independent methods and by a variety of different laboratories, as well as novel estrogen associated gene expression programs. These data indicate that future studies with flow-enriched genomics, transcriptomics, and proteomics will be able to identify cell-type specific mechanisms of estrogen action.

While the fixed-tissue analyses give information about heterogeneity of estrogen response, they have inherently low temporal resolution due to the need for sacrifice and the cross sectional nature of the experimental design. To record neuro-seeER longitudinally in the same populations of hypothalamic neurons in behaving animals, we virally transduced the MPO+MPN and VMH+TUof female Esr1*-*Cre mice, and implanted them with 200μm fibers over the MPO and ventrolateral VMH (VMHvl) (Fig. 3A, C; Fig. 5C). Following OVX (Fig. 3E), we imaged longitudinally across long time periods in the home cage using a custom, light-weight patchcord^9^. To image over long time periods without photo-bleaching, we employed a “snapshot” photometry technique, which alternates pulses of 470 nm and 560 nm LED light with 5 min between pulse pairs (Fig. 3B, D). To test whether neuro-seeER responds to exogenous E2, we recorded the neuro-seeER response in the same animal following injection of a vehicle control (oil, s.c.) and subsequently an injection of E2 (40 μg, s.c.). Both sets of injections were initiated at the same timepoint in the individual’s dark cycle with MPO and VMHvl neuro-seeER dynamics recorded simultaneously (Fig. 3F-G). To validate that the response of the EGFP is dynamic over time, we binned the fluorescence into h-long windows and compared how the signals changed across the snapshot recording session (Fig. 3H-K). Across timepoints, we observe that the neuro-seeER signal deviates significantly from the baseline pre-injection following the E2 injection (yellow: MPO; green: VMHvl) relative to oil control (black). Specifically, in both MPO and VMHvl, EGFP fluorescence was greater than baseline levels by 4 h post-treatment, and these elevated levels were maintained through 6 h and 16 h post-treatment (Fig. 3H). These data confirm our results showing estrogen responsive neurons in the MPO and extend those findings to the VMHvl.

**Fig. 3:**
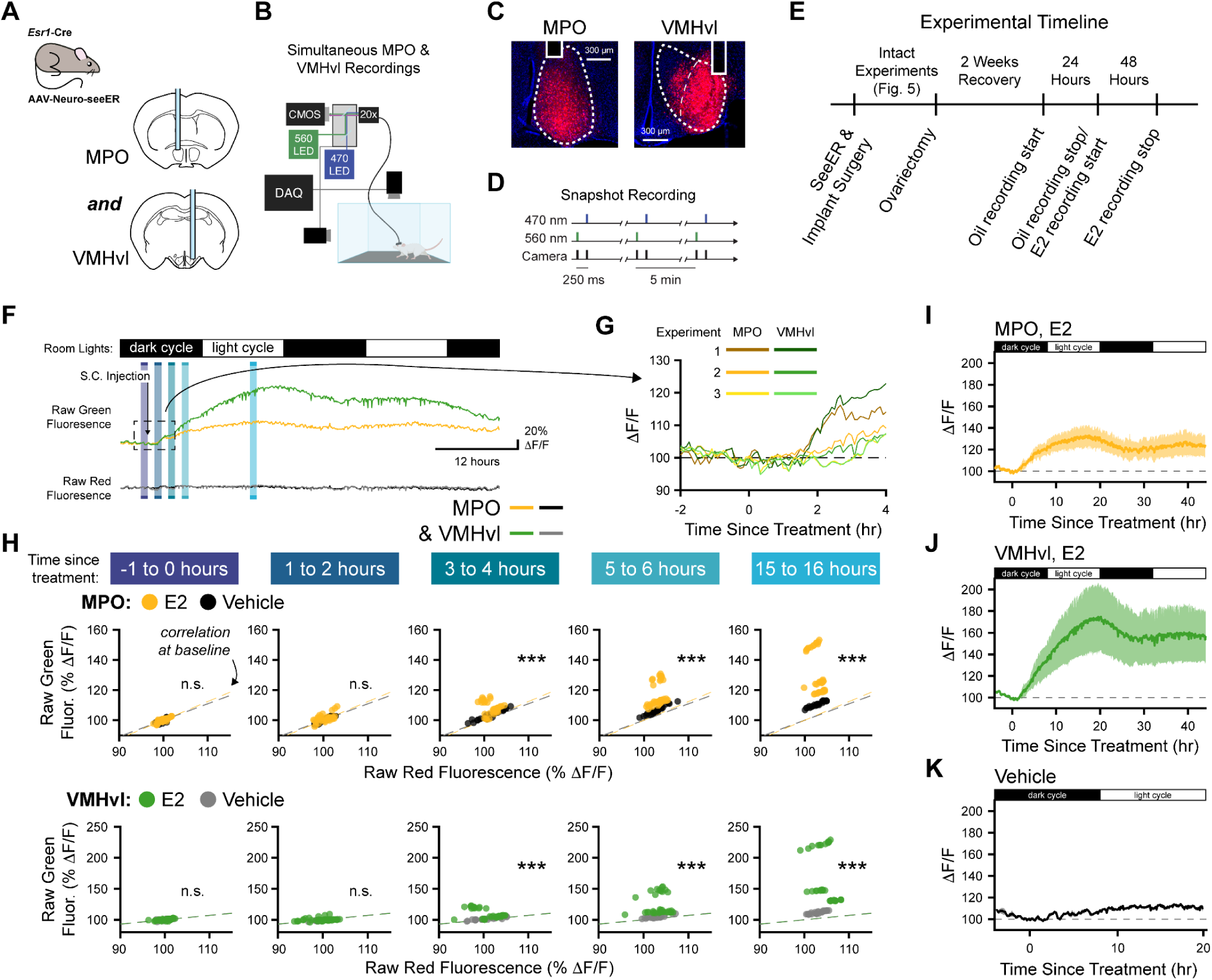
neuro-seeER validation *in vivo*. A, Schematic outlining viral strategy. B, Schematic depicting experimental setup for simultaneous 2-color neuro-seeER imaging in VMH. C, Example histology. D, Schematic illustrating Snapshot Recording technique where 470 nm and 560 nm pulse are given once every 5 minutes. E, Experimental timeline. F, Example of simultaneously recorded MPO+MPN (yellow) and VMHvl (green) neuro-seeER signals in response to an injection of 40 μg E2. Dashed rectangle indicates time window shown in more detail in H. Shaded rectangles indicate analysis windows used in i. Arrow: subcutaneous E2 injection. G, Detailed view of neuro-seeER response onset in simultaneously recorded MPO+MPN and VMHvl (N = 3 experiments, 2 mice). H, Raw green and red neuro-seeER fluorescence at -1 to 0, 1 to 2, 3 to 4, 5 to 6, and 15 to 16 h post injection in simultaneously recorded MPO+MPN and VMHvl populations (\*\*\**p* < 0.001, Linear Mixed Effects Model, N = 12 data points per experiment per time window. N_E2_ = 3 experiments, 2 mice; N_Veh_ = 2 experiments, 2 mice). I, Mean MPO+MPN neuro-seeER response to 40 μg E2 (N = 3 experiments, 2 mice). J, Mean VMHvl neuro-seeER response to 40 μg E2 (N = 3 trials, 2 mice). K, Mean MPO+MPN and VMHvl neuro-seeER responses to oil vehicle (N = 4 trials, 2 mice). See Table S1 for Linear Mixed Effects Model details.

### Neuro-seeER expression dynamically responds to endogenous fluctuations across the estrous cycle

To reveal dynamics of endogenously fluctuating estrogens, we transduced neuro-seeER by stereotaxic microinjection to two estrogen receptor expressing regions of the hypothalamus, the VMH+TU and MPN+MPO (Fig. 4A). Mice were tracked by vaginal cytology and sacrificed in each of the four estrous cycle stages to quantify estrogen responsiveness of the local neural populations. Simultaneously, a cohort of mice that did not express neuro-seeER was similarly tracked and sacrificed, followed by LC-MS/MS profiling of ovarian, serum, and brain E2. In both experiments, uterine weight reflected high levels of E2 in proestrus, as predicted^34^ (Fig. 4B). No significant differences were observed in uterine weights between the experiments. LC-MS/MS revealed a significant effect of estrous cycle stage on ovarian levels of E2 (Fig 4C), as well as in circulation (Fig. 4D). In bilateral 1-mm biopsy punches taken from the regions encompassing the VMH and MPO+MPN, cycle stage was significantly predictive of E2 levels (Fig. 4E). Comparison of E2 levels between hypothalamic nuclei revealed lower estrogen concentrations in the VMH+TU compared to MPO+MPN (*p* < 0.05) without any significant interaction between cycle stage and hypothalamic nucleus. Estrone (E1), 17α-estradiol, estriol (E3), and estetrol (E4) were detectable in the ovary (Fig. S3A-D) but not detected (N.D.) in the serum or brain. In humans, these N.D. estrogens play important roles in pregnant and postmenopausal, but not cycling, women^35^.

**Fig. 4.**
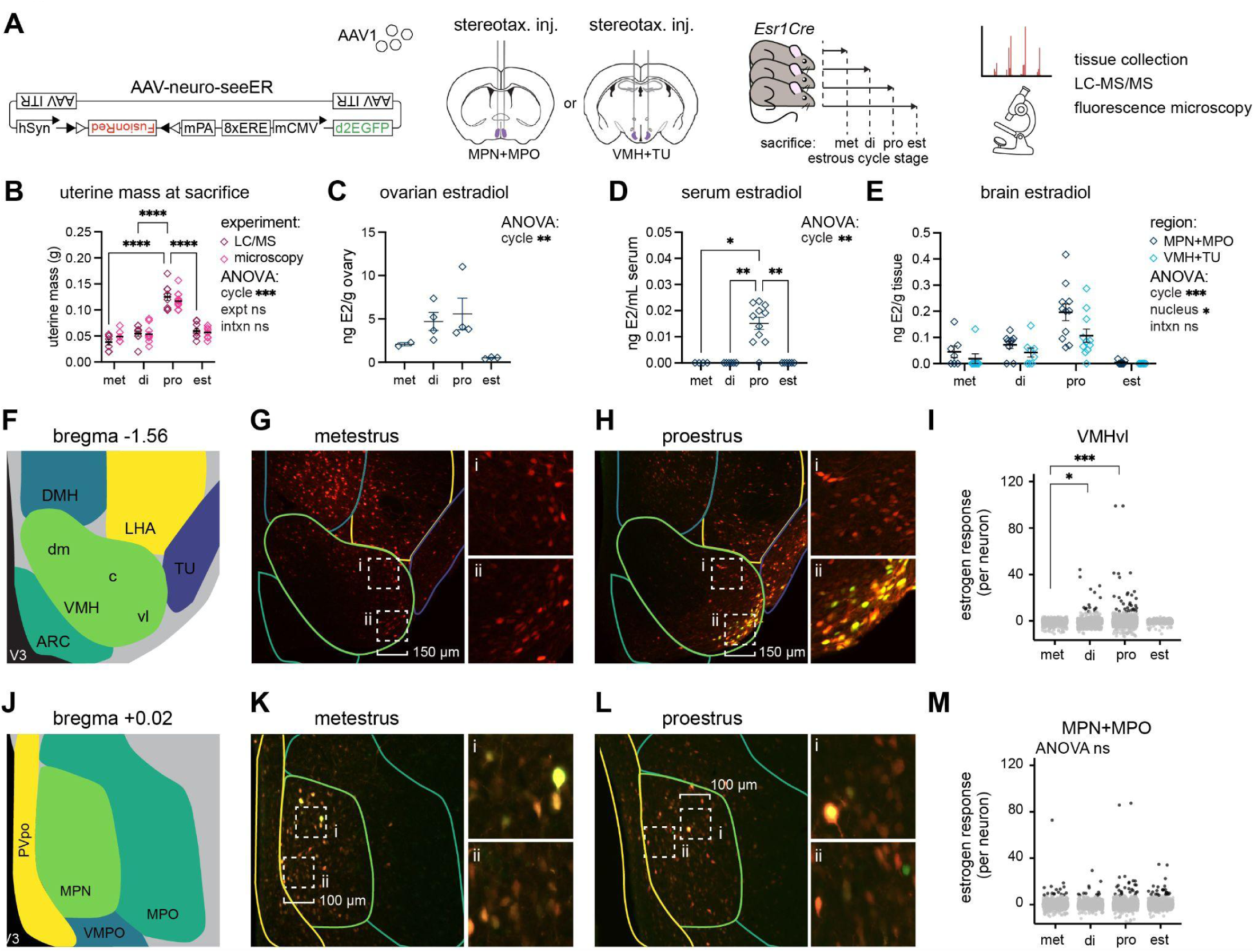
Estradiol levels and neuro-seeER expression across the estrous cycle. A, AAV harboring the neuro-seeER was delivered bilaterally to two regions of adult Esr1-Cre mice and analyzed across the estrous cycle (met = metestrus, di = diestrus, pro = proestrus, est = estrus). Separate cohorts were sacrificed for either liquid chromatography with tandem mass spectrometry (LC-MS/MS) based hormone quantification or fluorescence microscopy. B, Uterine mass (individual values and mean) across the estrous cycle for mice analyzed in each experiment. C, Ovarian E2 levels across estrous cycle stages. D, Serum E2 levels across estrous cycle stages. E, Brain E2 levels across estrous cycle stages taken from punches encompassing either VMH+TU or MPN+MPO. F, Anatomical boundaries of the VMH and surrounding regions analyzed at approximate Bregma -1.56 mm. G-H, FusionRed and EGFP expression in the VMH and surrounding regions in metestrus (G) and proestrus (H). I, Magnitude of estrogen response in non-responsive (gray) and responsive (green) neurons in the ventrolateral VMH across the estrous cycle. J, Anatomical boundaries of the MPN+MPO analyzed at approximate Bregma +0.02 mm. K-L, FusionRed and EGFP expression in metestrus (K) and proestrus (L). M, Magnitude of estrogen response in non-responsive (gray) and responsive (green) neurons in the MPN+MPO. Abbreviations: MPN, medial preoptic nucleus, MPO, medial preoptic area, PVpo, periventricular hypothalamic nucleus, preoptic part, ARC, arcuate hypothalamic nucleus, DMH, dorsomedial nucleus of the hypothalamus, LHA, lateral hypothalamic area, VMH, ventromedial hypothalamic nucleus; c = central part, dm = dorsomedial part, vl = ventrolateral part), V3, third ventricle, TN, tuberal nucleus of the hypothalamus. met: metestrus, di: diestrus, pro: proestrus, est: estrus. *, p<0.05; ***, p<0.001, ****, p<0.0001. See Table S1 for N’s, ANOVA details, and expanded pairwise comparisons.

We compared E2 levels with neuro-seeER fluorescence in the VMHvl (Fig. 4F) and found that both the levels of E2, and the number of estrogen responsive/EGFP-positive neurons (Fig. 4G-H) were significantly higher in diestrus and proestrus than in metestrus (Fig. 4I). In contrast to the VMHvl, we found that for the MPO+MPN (Fig. 4J), both E2 and the number of estrogen responsive neurons were highest in proestrus but only the hormone levels varied significantly across the cycle (Fig. 4J-M), indicating site-specific variability, despite similar cytology staging. The number of estrogen responsive neurons did not vary with estrous cycle stage in dorsomedial and central VMH (VMHdm+VMHc, Fig. S3E,F). These results reveal that, despite higher levels of circulating and brain-resident E2 during proestrous, there remains unexplored heterogeneity in the responsivity of individual neurons across the hypothalamus and even within individual hypothalamic nuclei. Surprisingly, we found that EGFP fluorescence varied significantly across the estrous cycle for VMH+TU but MPO+MPN neurons, despite higher local estrogen concentrations in MPO+MPN. These findings highlight the insufficiency of measuring E2 levels alone as a proxy for their effect on neural populations.

Finally, we tested whether neuro-seeER could be used *in vivo* to track endogenous fluctuations in estrogen response in the hypothalamus across long timescales. By providing temporally resolved neural population recordings of neuro-seeER across the estrous cycle, this approach allows us to directly test temporal heterogeneity of neural responses to fluctuating estrogens within the same individual.

We expressed neuro-seeER in the VMH and MPO+MPN of female Esr1-Cre mice, and implanted fibers over the VMHvl and MPO (Fig. 5A, C). Using our “snapshot” recording technique (Fig. 3B), we recorded neuro-seeER signal dynamics simultaneously in the VMHvl and MPO across an estrous cycle in female mice (Fig. 5B, D). We found a significant increase in neuro-seeER signal in the VMHvl during diestrus and proestrus and in the MPO during diestrus relative to that population’s neuro-seeER response during metestrus (Fig. 5E). Consistent with this, VMHvl neuro-seeER signal magnitude peaked later in the estrous cycle compared to the MPO signal (Fig. 5F) and was less variable. Despite clear estrous stage-dependent responses at the population level, neuro-seeER responses showed individual variability with individual experiments showing peak neuro-seeER fluorescence in diestrus, proestrus, or estrus (Fig. 5D). Strikingly, simultaneously recorded VMHvl and MPO neuro-seeER dynamics correlate with each other (Fig. 5H, “matched”), suggesting both populations are responding to the same endogenous fluctuation in local estrogens. To test this directly, we compared signal correlations when the signals were aligned to the estrous cycle (Fig. 5D) or aligned to peak neuro-seeER response (Fig. 5G). We found greater correlations between matched signals compared to mismatched signals only when the recordings were aligned to estrous (Fig. 5H). Together, these data indicate that neural responses to endogenous estrogen fluctuation show sequential, yet similar, dynamics in both recorded hypothalamic populations, and that there is notable individual variability in the peak estrogen response relative to estrous stage as estimated from vaginal cytology.

**Fig. 5.**
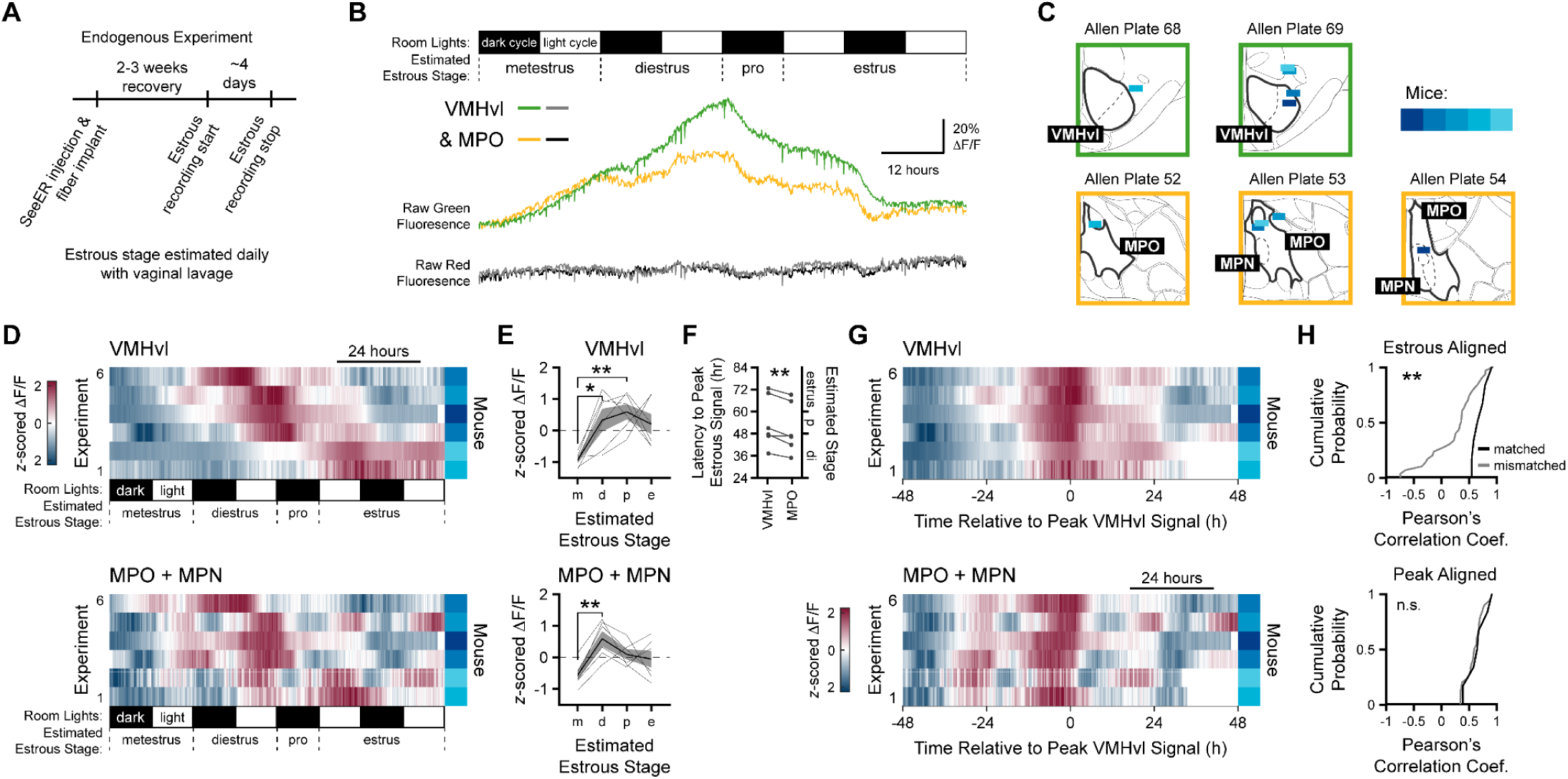
*In vivo* neuro-seeER imaging reveals spatiotemporal differences in the brain’s transcriptional response across the estrous cycle. A, Timeline for endogenous E2 experiments. Estrous stage was estimated using epithelial cytology obtained daily from vaginal lavage. B, Example of simultaneously recorded MPO+MPN (yellow) and VMHvl (green) neuro-seeER signals across an estrus cycle. Estimated estrous stages shown above data. C, MPO+MPN (yellow boxes) and VMHvl (green boxes) histology. Mouse identity is color coded in blue and is matched across MPO+MPN and VMHvl. D, Heatmaps of simultaneously recorded MPO+MPN (top) and VMHvl (bottom) neuro-seeER signals across an estrus cycle. Experiments are sorted by timing of the peak of the VMHvl neuro-seeER signal. Estimated estrous stage shown below heatmaps. N = 6 experiments, 5 mice. E, Mean neuro-seeER signal in MPO+MPN (top) and VMHvl (bottom) across estimated estrous stages (m: metestrus, d: diestrus, p: proestrus, e: estrus). MPO+MPN: Main Effect of Estrous Stage, F_3, 15_ = 3.86, *p* = 0.031. VMHvl: Main Effect of Estrous Stage, F_3, 15_ = 4.90, *p* = 0.014, 1-way RM ANOVAs. N = 6 experiments, 5 mice. Post hoc tests: \**p* < 0.05, \*\**p* < 0.01. Tukey-Kramer method used for post hoc statistics. Data displayed as mean ± S.E.M. F, Latency of peak estrous signal relative to metestrus onset. Effect of brain region: t_5_ = 4.13, *p* = 0.0091. G, Heatmaps of simultaneously recorded MPO+MPN (top) and VMHvl (bottom) neuro-seeER signals across four days aligned to peak neuro-seeER response. Experiments are sorted as in *D*. Time relative to peak signal shown below heatmaps. N = 6 experiments, 5 mice. H, Top: Cumulative distribution of MPO+MPN/VMHvl correlations from simultaneous recordings (matched, black) or recorded separately (mismatched, gray). Two-sample Kolmogorov-Smirnov test, D_35_ = 0.7667, *p* = 0.0022. Bottom: Same as top, but signals are aligned to the peak neuro-seeER response. Two-sample Kolmogorov-Smirnov test, D_35_ = 0.2667, *p* = 0.81. N = 6 matched comparisons, 30 mismatched comparisons, 6 experiments, 5 mice. See Table S1 for ANOVA details.

## Discussion

For decades, immunodetection of estrogen receptors has served as the gold standard for defining estrogen-sensitive tissues and cell types. Yet estrogen signaling is inherently dynamic, and static measures of receptor presence fail to capture the dynamic effects of fluctuating estrogen levels and heterogeneous responses across tissues and cell types. Here, we introduce neuro-seeER, a bifunctional reporter that distinguishes estrogen sensitivity from functional estrogen-induced responses in the brain with cell-level resolution, and we demonstrate its utility *in vivo* to track changes in exogenous and endogenous estrogen changes. Neuro-seeER is built on the framework of earlier ERE-luciferase and ERE-GFP systems, expanding their utility by enabling direct visualization of dynamic estrogen responses in individual neurons across multiple hypothalamic nuclei and estrogen fluctuations across the estrous cycle. This approach fills an important technical gap and demonstrates how longitudinal tracking of estrogen responsivity can reveal unexpected heterogeneity at the cellular and individual levels.

*In vitro*, seeER EGFP expression was dynamic, ligand-dependent, and receptor-specific. As expected, both of the classical nuclear estrogen receptors, ERα and ERβ, induced an increase in the number of EGFP+ cells in a ligand dependent fashion, whereas the evolutionarily similar nuclear receptors, AR and GR, showed no induction of EGFP+ cells in response to E2. These results confirm that the seeER produces EGFP only when both an ER and an estrogen are present, making it a faithful and specific reporter of estrogen responses. *In vivo*, the neuro-seeER was induced by endogenous and exogenous estrogens, as observed in cross sectional microscopy experiments, and in within-subject longitudinal, multi-site fiber-photometry imaging.

By nature, estrogen responses are multifaceted and cannot be captured with a single metric. While we demonstrate the utility of using neuro-seeER for tracking estrogen response across a wide variety of neuroendocrinology experiments, several critical caveats prevent it from detecting all potential estrogen responses. First, because of technical limitations, some aspects of the estrogen response may be missed by our current approach. The neuro-seeER construct uses 8 repetitions of the consensus ^36^ ERE motif, the most commonly found sequence associated with ERα binding in the brain ^37^. However, estrogen receptors can also bind to slight modifications of the consensus ERE with reduced affinity^36^ and the current neuro-seeER does not capture these dynamics.

Future customized versions of seeER may be able to better detect these low-affinity interactions. Secondly, neuro-seeER is a transcriptional reporter and thus only captures the genomic or “classic” actions of ERs. The neuro-seeER cannot report the “non-classical” effects of membrane-associated estrogen receptors, such as GPER1^38^ as well as membrane-localized ERα^39,40^ and ERβ^41^. These limitations also represent fruitful avenues for future study and tool development. Nonetheless, the canonical nuclear activities of ERα are critical for reproductive physiology^42^ and the development of breast cancers^43^. In addition, the robust EB, E2, and cycle-dependent responses observed here demonstrate the utility of the neuro-seeER for visualizing and localizing dynamic estrogen responses. Future experiments can multiplex imaging of longitudinal slow genomic estrogen signatures with fast non-genomic responses using modified estrogen detecting GPCRs, which will allow an even fuller picture of estrogen responsivity to emerge.

*In vivo,* the neuro-seeER uncovered unexpected heterogeneity in estrogen responsiveness. Within each estrogen-sensitive nucleus examined (MPO, VMH, DMH), only subsets of neurons engage robust ERE-driven transcription, while neighboring neurons remain relatively unresponsive. In the VMH, these high-response neurons were confined to the ventrolateral subdivision and were only observed in diestrus and proestrus, when E2 levels peak. High response in proestrus is consistent with the VMHvl’s established role in regulating female sexual behavior in a cycle-dependent manner^44^. In addition, we found reduced responsiveness of MPO compared to VMHvl neurons across the estrous cycle, despite greater concentrations of E2 in the MPO. The observed dynamic heterogeneity suggests that responsiveness can be coupled, or uncoupled, to circulating and local hormone levels depending on unknown factors. Additional regulatory mechanisms that determine whether an estrogen receptor expressing cell responds to the presence of hormone may include expression of transcriptional coactivators and corepressors^45^, post-translational modifications to key proteins^46^, intracellular sequestration^47^, and/or changes to the chromatin landscape of regulated genes^48^. Dynamic heterogeneity in estrogen response also aligns with a broader principle of hypothalamic organization: molecular and functional heterogeneity even within individual hypothalamic nuclei^21,49–51^. Just as hypothalamic nuclei are composed of neural populations that are distinct, specialized, and intermingled, estrogen-sensitive neurons comprise a mosaic of responsive states. By exposing this diversity, neuro-seeER reframes estrogen signaling not as a uniform property of ERα+ neurons but as a process that is dynamic beyond the typical steroid fluctuations seen in the estrous cycle.

This framework carries broad implications: it suggests that changes in estrogen responsiveness, rather than receptor expression *per se*, may underlie hormone-dependent changes in reproduction, thermoregulation, metabolism, and social behavior. In addition, our work here establishes a generalizable strategy for studying hormone responsiveness. The neuro-seeER design principle has broad applicability: it can be adapted to glial populations, other nuclear hormone receptors, and/or other tissue types (*e.g.*, uterus, breast tissue, bone) to examine the dynamic heterogeneity of cellular responses to steroid hormones. Estrogen response detection *in vivo* can be achieved using optical off-the shelf methods, such as fiber photometry or microendoscopy, allowing new integration with population or cellular resolution recordings of neural activity. Together, neuro-seeER initiates a new paradigm for separating estrogen sensitivity from estrogen responsiveness, reveals unappreciated cellular and regional diversity in hormone signaling, and opens new avenues for dissecting how steroid hormones dynamically shape neural circuits, neural computation, and behavior.

## Methods

### Tissue culture

293T cells were freshly purchased from ATCC (Cat no: CRL-3216). 293T cells were seeded to 50,000 cells per well in 24-well plates. Cells were maintained in phenol red-free DMEM/High Modified media (Cytiva SH30284, Marlborough, MA) supplemented with penicillin, streptomycin, and charcoal-stripped fetal bovine serum (Gibco A56707, lot #U2979526RP). 24 h after seeding, receptor and reporter DNA were transfected in a 1:3 ratio using the ViaFect Transfection Reagent (Promega E4981, Madison, WI) per supplier’s recommendations. Following another 24 h, hormone (17β-estradiol (E2) or 5α-dihydrotestosterone(DHT)) suspended in molecular grade ethanol, or vehicle, was added to the cells. After hormone or vehicle administration, the cells were placed in an Incucyte SX5 (Sartorius, Gӧttingen, Germany). Cells were imaged every 2 h across a 96 h period at 10X magnification. After each imaging session, Incucyte software was used to count green objects.

### Animals

Esr1-Cre mice^52^ (*Esr1^Cre^*, B6N.129S6(Cg)-*Esr1^tm1.1(cre)And^*/J , JAX stock # 017911) were maintained on a C57BL/6J or CD1 background and housed in Association for Assessment and Accreditation of Laboratory Animal Care (AAALAC) approved facilities at the University of California, Los Angeles and Princeton University in accordance with recommendations in the Guide for the Care and Use of Laboratory Animals and the UCLA Institutional Animal Care and Use Committee (IACUC). All studies were carried out with IACUC approval. At UCLA, mice were maintained under a 12:12 h light/dark cycle (lights on at 7:00 AM PST, Zeitgeber time (ZT) = 0), with ambient temperatures ∼22°C, and 30-70% relative humidity. At Princeton, mice were maintained under a 12:12 h reversed dark:light cycle (lights off at 10:00 AM EST, Zeitgeber time (ZT) = 12), with ambient temperatures 21-26°C, and 30-70% relative humidity. All mice were provided water and food ad libitum. Female mice were defined as animals with a relatively shorter anogenital distance, mammary glands, and the presence of ovaries confirmed post-mortem.

### Reporter Constructs

A dual-function adeno-associated virus (AAV) vector was engineered with a flip-excision (FLEX^53^) switch to drive Cre-dependent expression of a red fluorescent protein (FusionRed) under the control of a human synapsin 1 (hSyn) promoter reported to give highly specific expression in neurons (PMID: 12595892). Outside of the FLEX cassette, eight tandem EREs (8xERE) drive expression of a destabilized enhanced green fluorescent protein (d2EGFP) under a minimal mouse cytomegalovirus (mCMV) promoter (Fig. 1A). The destabilized EGFP has a reported 2 h (h) half-life which confers temporal specificity^26^. Presence of Cre-dependent FusionRed allows for identification of *Esr1*+ (which encodes ERα) cells, confirms vector expression independent of estrogen response, and is restricted to neuronal expression via the hSyn promoter. FusionRed was chosen as the red fluorescent protein due to minimal cytotoxicity to mammalian cells^54^ . The neuro-seeER virus (AAV-hSyn-FLEX-FusionRed-mCMV-8xERE-d2EGFP, serotype 1) was generated by the University of North Carolina Vector Core, with a titer of 3.5x10^12^ virus molecules/mL. A ubiquitously expressed reporter construct (CMV-FLEX-FusionRed-mCMV-8xERE-d2EGFP) was generated for transfecting into cells for *in vitro* validation studies.

### Custom Multisite Photometry

Photometry was performed using a customizable multisite fiber photometry platform. Details on multisite implant and patchcord fabrication can be found in our prior publications^9,55^. Briefly, these designs allow for customizable photometry to target multiple defined stereotactic coordinates using 200 μm optic fibers.

### Stereotaxic Surgery

Brain regions targeted for injection were chosen based on their abundance of ERα expression ^56,57^ and previous studies demonstrating roles in regulation of temperature, metabolism, and/or reproduction^58–60^ which are of particular interest to our lab. Target regions included the MPN, which is surrounded by the MPO, and the ventrolateral portion of the ventromedial hypothalamus (VMHvl), which is adjacent to the dorsomedial hypothalamus (DMH) and tuberal nucleus (TU)^61^. For all experiments except longitudinal photometry, adult Esr1-Cre mice (10-13 weeks of age) were anesthetized with isoflurane, and AAVs were injected bilaterally with a glass pipette (500 nL per side) to the MPO (coordinates from Bregma: anterior-posterior, +0.20 mm, medial-lateral, +/- 0.35 mm, dorsal-ventral, -5.3 mm), and/or VMH (coordinates from Bregma: anterior-posterior, -1.5 mm, medial-lateral, +/- 0.85 mm, dorsal-ventral, -5.5 mm) as previously described (41). For photometry experiments, adult mice (12-24 weeks of age) were anesthetized (isoflurane: 3-5% induction; 1-2% for maintenance), and mouse skulls were leveled using confluence of the sagittal sinus and rostral rhinal vein (lateral vein that sits on the dorsal surface of the mouse brain between the olfactory bulb and prefrontal cortices) as the anterior position and lambda as the posterior position. AAV injections (300 nL/site; coordinates from the confluence of the sagittal sinus and rhostral rhinal vein; dorsal-ventral distance measured from brain surface) targeted the MPO (anterior-posterior, -3.00 mm, medial-lateral, - 0.40 mm, dorsal-ventral, -5.0 mm) and VMHvl (anterior-posterior, -4.65 mm, medial-lateral, +0.72 mm, dorsal-ventral, -5.8 mm). Subsequently, a custom multisite photometry probe was implanted at fiber depths of -4.55 mm at the MPO and -5.5 at the VMHvl as described previously. At UCLA, mice were provided buprenorphine and carprofen for recovery and monitored daily for one week. At Princeton, mice were provided ketofen for recovery and monitored daily for one week.

### Estrous Cycling and Ovariectomy

Starting at least one week following stereotaxic surgery (STX), vaginal lavage was performed daily (between ZT +3 to +4 or ZT +15 to +16) to assess estrous cycle stage based on the proportion of leukocytes, nucleated epithelial, and keratinized/cornified epithelial cells ^62^. At least two complete estrous cycles were evaluated prior to sacrifice. For non-photometry experiments, mice were sacrificed during proestrus (P, n = 4-6), estrus (E, n = 3-5), metestrus (M, n = 3-4), or diestrus (D, n = 3-7), between ZT +4 to +6. Longitudinal imaging mice were ovariectomized (OVX) following the completion of estrous photometry experiments. Photometry mice were provided buprenorphine and ketofen for recovery, were monitored daily for one week, and were allowed to recover for at least two weeks prior to beginning post-OVX imaging experiments. Photometry mice were sacrificed following the completion of post-OVX experiments. A separate cohort of mice received were ovariectomized (OVX) one week following STX, allowed to recover for one week, then injected with 1 μg of estradiol benzoate (EB, β-estradiol 3-benzoate, Sigma-Aldrich, Cat # E8515) in 100μL corn oil + 0.1% DMSO, and sacrificed after 4, 6, or 16 h (n =4/group). Control OVX mice were injected with vehicle only (100μL corn oil + 0.1% DMSO, n = 6) and sacrificed after 4, 6, or 16 h.

### Tissue Collection and Preparation

Prior to tissue collection, all mice were deeply anesthetized with sodium pentobarbital. For acute experiments, the uterus was dissected and weighed. Uterine wet mass was used to confirm estrous cycle stage at time of sacrifice (uterine mass >100 mg for proestrus^63^). Blood was collected from the inferior vena cava, allowed to clot for 90 min at room temperature, then centrifuged for 15 min at 2,000 x g. Trans-cardial perfusion was performed with 1x phosphate buffered saline (PBS) until liver and lungs cleared (∼1-2 min), followed by ice cold 4% paraformaldehyde (PFA) in 1x PBS for 10 min. Brains were dissected and post-fixed in 30% sucrose + 4% PFA in 1x PBS for 24 h, then transferred to 30% sucrose in 1x PBS (24 h) until embedding in optimal cutting temperature (OCT)-compound and storage at -80°C. For animals with multisite photometry implants, skulls with implants intact were allowed to post-fix in 4% PFA in 1x PBS for at least 3 days before brain dissection to ensure visible fiber tracts following cryosectioning. Brains were sectioned at 20-50 µm thickness on the coronal plane into six series on a Leica CM1950 cryostat. Sections were either collected directly onto SuperFrost Plus or Excell Slides, or collected free floating in 1x-PBS and stored in cryoprotectant at -20°C (30% sucrose, 30% ethylene glycol, 1% polyvinylpyrrolidone in 1x PBS).

### Immunohistochemistry

Brain sections were briefly rinsed with 1x PBS, then blocked for 1 h with 5% normal goat serum with 0.2% Triton X-100 in 1x PBS at room temperature. Sections were incubated overnight at room temperature with rabbit anti-estrogen receptor α (ERα, 1:5000, Millipore Cat # 06-935, RRID: AB_310305) antibodies. Sections were rinsed with 1x PBS and then incubated with goat anti-rabbit AlexaFluor 647 antibodies (1:500, Jackson ImmunoResearch Cat # 111-606-003, RRID: AB_2338079) for 1 h at room temperature. For reference ERα expression, a separate set of sections from wild-type female mice were dehydrated in graded ethanol, dried, then heated to 95°C for 40 min in antigen retrieval buffer (25 mM Tris-HCl, pH 8.5, 1 mM EDTA, 0.05% SDS).

Sections were then blocked for 1 h with 5% normal goat serum with 0.2% Triton X-100 in 1x PBS at room temperature, then incubated with mouse anti-ERα antibodies (1:250, Santa Cruz Cat # sc-8005, RRID: AB_627556) overnight at room temperature. Sections were rinsed with 1x PBS and then incubated with goat anti-mouse AlexaFluor 647 antibodies (1:500, Jackson ImmunoResearch Cat # 115-606-003, RRID: AB_2338921) for 2 h at room temperature. Sections were counterstained with DAPI (1:1000) for 30-60 sec. Slides were coverslipped with Fluoromount-G mounting medium (ThermoFisher Scientific). For brains from mice receiving photometry implants, slices were washed in PBS and mounted on slides, allowed to dry, coverslipped with a mounting medium (EMS Immuno Mount DAPI and DABSCO, Electron Microscopy Sciences, Pennsylvania, USA), and clear nail polish was used to seal the coverslip to the slide. FusionRed and EGFP did not require antibody amplification.

### Steroid Measurement

To determine if the specific levels and steroid profile of brain nuclei were correlated with ERE reporter expression, we measured levels of estrogens, including E2, in the brain regions of interest (MPN/MPO and VMH/TU). A separate cohort of adult (10-19 weeks of age) wild-type female mice (Esr1-Cre wild-type littermates or C57BL/6J, JAX #000664, RRID: IMSR_JAX:00064), were estrous cycle staged as described above. During proestrus, estrus, metestrus, or diestrus, female mice were deeply anesthetized with isoflurane, then rapidly decapitated (proestrus n= 11, estrus n = 9, metestrus n = 7, diestrus n = 9). Trunk blood was collected and processed for serum as described above. Fresh brains were rapidly embedded in OCT compound and snap frozen on dry ice with absolute ethanol. No more than 3 min elapsed between anesthetic induction and embedding. Uterine wet mass was recorded to confirm cycle stage. One ovary was weighed, then frozen on dry ice and stored at -80°C.

### Brain Microdissection

Fresh frozen brains in OCT were thawed to -12°C for at least 1 h prior to sectioning on a Leica CM1950 cryostat. Brains were sectioned at 300 µm on the coronal plane, and were mounted onto a Superfrost Plus slide. Sections were allowed to chill to -20°C before microdissection. A chilled stainless-steel biopsy punch tool (Fine Science Tools, Cat # FST-18035-01, inner diameter 1 mm, tissue wet weight mean 0.24 mg/punch) was used to collect four tissue punches each (two per side over two serial sections) for the MPO approximately at the level of the MPN and a mediobasal region that includes the VMH, DMH, and TU. A separate punch tool was used for each nucleus and each experimental group. Tissue punches were collected into chilled 2 mL polypropylene tubes (Sarstedt, Cat # 72.694.007) containing five 1.4 mm zirconium ceramic oxide beads (Fisher Scientific, FisherBrand Cat #15-340-159), and stored at -80°C until processing.

### Steroid Extraction

Steroids were measured by liquid chromatography tandem mass spectrometry (LC-MS/MS) in ovary, serum, and two microdissected brain regions (MPN with surrounding MPO and VMHvl with adjacent TU). The analytes included estrone (E1), E2, 17α-estradiol (17α-E2), estriol (E3), and estetrol (E4). Serum and brain samples, but not ovary samples, were derivatized using 1,2-dimethylimidazole-5-sulfonyl-chloride (DMIS)^64^, which increases sensitivity for estrogens by approximately 10-fold^65^.

A subset of total collected ovary samples (proestrus n = 4, estrus n = 3, metestrus n = 2, diestrus n = 4) were used as a positive control to ensure successful detection of target steroids prior to measurement of serum and brain samples. Frozen ovaries were weighed and placed in 2 mL polypropylene tubes as described above. Ovaries were homogenized in 1 mL of HPLC-grade acetonitrile for 30 sec using a bead mill homogenizer. A volume of homogenate equivalent to 1 mg of ovary was used for steroid extraction. Ovary homogenates were processed using liquid-liquid extraction^66,67^. In brief, we added 50 μL total of ^13^C labelled internal standards (^13^C-E2 and ^13^C-E3, Cambridge Isotope Laboratories) in 50:50 HPLC-grade methanol:MilliQ water to each sample except “double blanks.” Then 1 mL of HPLC-grade acetonitrile was added, samples were vortexed, and samples were centrifuged at 16,100 *g* for 5 min. Then 1 mL of supernatant was transferred to pre-cleaned borosilicate glass culture tubes. Next, 0.5 mL of HPLC-grade hexane was added, and samples were vortexed and then centrifuged at 3,200 *g* for 2 min. The hexane was discarded, and the acetonitrile was dried in a vacuum centrifuge at 60°C for 50 min. Residues were resuspended in 55 μL of 25% HPLC-grade methanol in MilliQ water, transferred to 0.6-mL polypropylene microcentrifuge tubes, and centrifuged at 16,100 *g* for 2 min. Then 50 μL of supernatant were transferred to glass inserts in LC vials and stored at -20°C until injection for LC-MS/MS analysis. All samples were extracted alongside calibration curves, blanks, double blanks, and quality control (QC) samples. Calibration curves for all analytes ranged from 0.01 to 10,000 pg per tube.

Serum (20 μL), MPO/MPN (0.94 mg) and VMH/TU (0.705 mg) samples were processed by liquid-liquid extraction followed by DMIS derivatization^65^. During liquid-liquid extraction, instead of being resuspended in 25% methanol after drying, residues were placed in a wet ice bath and resuspended in 30 μL of sodium bicarbonate buffer (50 mM, pH 10.5). Then 20 μL of 1 mg/mL DMIS in acetone was added. Samples were vortexed, centrifuged at 3200 *g* for 1 min, and transferred to glass inserts in LC vials. Vials were capped and placed in a water bath at 60°C for 15 min. Samples were cooled at 4°C for 15 min, centrifuged at 3200 *g* for 1 min, and stored at -20°C until injection for LC-MS/MS analysis. Underivatized standards were included to measure any underivatized estrogens in samples that underwent derivatization. These standards received 20 μL of acetone without DMIS after resuspension in sodium bicarbonate buffer. In all derivatized standards, blanks, quality controls, and samples, we were unable to detect underivatized estrogens, demonstrating a reaction efficiency of 100%.

### Steroid Analysis by LC-MS/MS

Steroids were quantified using a Sciex QTRAP 6500 UHPLC-MS/MS system^65,67^. Samples were transferred to a refrigerated autosampler (15°C). For ovary samples, 45 μL were injected into a Shimadzu Nexera X2 UHPLC system, whereas for serum and brain samples, 35 μL were injected. Samples were passed through a KrudKatcher ULTRA HPLC In-Line Filter (Phenomenex, Torrance, CA, USA), then through a SecurityGuard^TM^ ULTRA C18 guard column (2.1 mm) (Phenomenex), and separated on a Kinetex® 2.6 µM EVO C18 100 A° LC column (Phenomenex; 2.1 50 mm; 2.6 μm; at 40°C) using 0.1 mM ammonium fluoride in MilliQ water as mobile phase A (MPA) and HPLC-grade methanol as mobile phase B (MPB). The flow rate was 0.4 mL/min. During loading, the gradient profile was at 10% MPB for 0.5 min. From 0.6 to 4 min, the gradient profile was increased to 42% MPB, then to 60% MPB until 9.4 min. From 9.4 to 9.5 min, the gradient was at 60-70% MPB and increased to 98% MPB until 11.9 min. Finally, a column wash was carried out from 11.9 to 13.4 min at 98% MPB. The MPB was then returned to the starting conditions of 10% MPB for 1 min. The total run time was 14.9 min. Before and after each sample injection, the needle was rinsed externally with 100% methanol.

Quantification was achieved through multiple reaction monitoring (MRM), with two MRM transitions for analytes and one MRM transition for internal standards. The MRMs were scheduled based on known retention times (180 sec windows). Analytes were ionized using electrospray ionization. Derivatized estrogens were ionized using positive mode, while underivatized estrogens were ionized using negative mode. The QCs at 5 different levels (0.5, 2, 20, 200, and 8000 pg) were prepared in neat solution and run in triplicate to determine assay accuracy and precision. Accuracy was assessed by comparing measured values with known values, and precision was assessed by the coefficient of variation (CV) across replicates. Accuracy of all QCs was acceptable (within 15%), and precision was acceptable (< 15%). All blanks and double blanks were non-detectable for analytes.

### Isolation of Hypothalamic Neurons for Flow Cytometry and RNA Sequencing

#### Isolation of Hypothalamic Neurons

One week following OVX, Esr1-Cre mice with the AAV-ERE-EGFP transduced bilaterally to the MPO and VMHvl were injected with 1 µg EB (n = 3) or vehicle (n = 3) subcutaneously as described above. Six h after injection, mice were deeply anesthetized with isoflurane, then rapidly decapitated. Fresh brains were quickly placed into a stainless-steel mouse brain matrix (Zivic Instruments, Cat # BSMAS001-1), and 1 mm coronal sections containing the preoptic area and mediobasal hypothalamus were isolated. Sections were placed into Hibernate A media (without calcium, BrainBits, Cat # HACA500), and with fluorescent illumination, FusionRed expressing regions were collected. Cells were then dissociated with papain (Worthington, Cat # LK003176) and DNase I (Worthington, Cat # LK003170) in Earl’s Balanced Salt Solution (Worthington, Cat# LK0031890) for 45 min at 37°C with gentle shaking. Cells were sorted in the UCLA BSRB flow cytometry core facility on a BD FACS ARIA II. The FACS gating pipeline was optimized to select living, nucleated cells that express the fluorophore FusionRed, followed by selection for high or low EGFP expression. Briefly, cell-sized objects were selected away from debris by forward scatter (FSC) and side scatter (SSC), followed by doublet discrimination. Live, nucleated cells were selected by DRAQ5 (ThermoFisher, Cat # 62251) positivity and DAPI (NucBlue, 4’,6-diamidino-2-phenylindole, ThermoFisher, Cat # R37606) negativity. Finally, cells were selected for FusionRed positivity based on the appearance of a clearly defined population not present in historical controls. Then EGFP was divided by relative intensity of green fluorescence. Bulk cells were sorted into nuclease-free 1x PBS, and centrifugated for 8 min at 600 x g at RT. Cells were then resuspended in 10 µL Trizol (Ambion, Cat # 15596018) and stored at -80°C until library preparation. Samples were selected for further processing if at least 450 cells in each population were recovered. A maximum of 3393 cells were submitted for library preparation. RNA extraction and cDNA library preparation for ultra-low input were performed by Azenta Life Sciences.

### Microscopy and Image Analysis

To confirm fiber targeting in photometry experiments, slides were imaged with a digital robotic slide scanner (NanoZoomer S60, C13210-01, Hamamatsu Photonics, Hamamatsu, Japan), and fiber tip locations were manually identified and localized using the Allen Mouse P56 Brain Atlas (atlas.brain-map.org, Washington, USA). For all other microscopy, digital images were acquired with a Nikon Eclipse Ti2 inverted microscope and Nikon NIS Elements AR. The same magnification, exposure, and illumination settings were used for images of the same experiment. Analysis was performed on one hemisphere of each animal at approximately the same rostro-caudal level of the MPN (approximate Bregma +0.02 mm), VMH (approximate Bregma -1.34 and -1.46 mm), and TU (approximate Bregma -1.46 mm). Brain nuclei boundaries and anatomical notations are in accordance with the Allen Mouse Brain Atlas (adult postnatal day 56; Coronal Reference Atlas, Allen Institute for Brain Science, Allen Mouse Brain Atlas; http://mouse.brain-map.org/static/atlas).

Unscaled tiff images were segmented using cellpose-SAM ^68^ and default parameters. Neuronal objects were identified using FusionRed images, and ImageJ compatible ROIs were saved. Quantification of FusionRed and EGFP fluorescence within each ROI was performed on unscaled microscopy images in Fiji(ImageJ) software ^69^. ROIs were grouped by hypothalamic nucleus using a custom ImageJ plugin allowing for user drawn hierarchical regions of interest. Together this pipeline allows for single-cell based quantification grouped by user-defined hypothalamic nucleus. All collected data were analyzed and visualized in R using the package tidyverse_2.0.0. Statistics were performed with a combination of published and custom packages: MASS_7.3-65, nlme_3.1-168, and microscopie_0.1.3. All packages are available on CRAN other than microscopie_0.1.3, which is available at https://github.com/jevanveen/microscopie/. To determine the “green bias” of individual cells, data were grouped by image in R, and a linear regression of green and red signal was performed. Residuals for green fluorescence were calculated using red signal as the independent variable. These residuals were then scaled and centered across all datasets using a robust z-score. A cutoff of z = 8 was chosen based on inspection of OVX+EB replacement data and applied to all *in vivo* analyses. Statistical testing to determine the effect of EB treatment or estrous stage was performed using a mixed effects negative binomial model ANOVA with individual mice treated as random effects. Reduced model tests were performed excluding treatment group as a factor, and a post-hoc ANOVAs were performed indicating in all cases that the full model significantly accounted for a greater percent of the variability in the data (p < .05). All data import, processing, graphing, and statistical analyses are included as supplemental information.

### Bioinformatics

Raw reads in fastq files were assembled to the genome using kallisto 0.46.2. Count tables were queried for differential gene expression using DESeq2 Galaxy Version 2.11.40.8+galaxy0. GSEAs were performed in R using the package ‘fgsea’ version 1.14.0. Graphs were made in R with ggplot2 3.5.2 and custom wrapper functions available in the package ‘grafiek’ at https://github.com/jevanveen/grafiek.

### Multifiber “Snapshot” Recording

Multisite fiber photometry recordings were made using an FP3002 (Neurophotometrics, California, USA). The FP3002 contains 470, and 560 nm LEDs for EGFP and FusionRed excitation, respectively. A dichroic mirror splits EGFP and FusionRed emission into separate channels where they are further filtered with bandpass filters (green channel: 494-531 nm; red channel: 586-627 nm). Mirrors reflect this emission onto distinct locations on the sensor of a CMOS camera. For “snapshot” recording of neuro-seeER fluorescence, 50 ms of 560 nm LED stimulation is presented, and is followed by 150 ms of no LEDs and then 50 ms of 470 nm LED stimulation. This pattern of LED stimulation was repeated every 5 minutes (0.00333 Hz) until the end of recordings. Custom Bonsai code controlled the FP3002 system.

### Data Processing

Data were saved by Bonsai as .csv files and were subsequently aligned to ZT and organized into dictionaries using custom Python code (Python Software Foundation, Delaware, USA). Raw neuro-seeER fluorescence, ZT timestamps, and injection time were then loaded into custom MATLAB code (Mathworks, Massachusetts, USA) for subsequent analysis. All analyses were performed on raw neuro-seeER fluorescence. Code used for photometry analysis is available at https://github.com/emguthman/neuro-seeer-manuscript.

### Photometry: Exogenous Hormone Treatment

Following OVX and recovery, neuro-seeER fluorescence was recorded using the FP3002 for 72 h. After an initial 4 h baseline, the animal was injected with 0.1 mL sesame oil vehicle control (Sigma-Aldrich, CAS #8008-74-0) and neuro-seeER fluorescence was recorded for a subsequent 20 h. Then, the recording was stopped, and a new one began. After an initial 4 h baseline, the animal was injected with 40 μg 17β-E_2_ (Sigma-Aldrich, CAS #50-28-2) dissolved in 0.1 mL sesame oil vehicle and neuro-seeER fluorescence was recorded for a subsequent 44 h. To compare the change in neuro-seeER fluorescence in response to 17β-E_2_ or vehicle, we quantified green and red channel fluorescence in response to 470 or 560 nm LEDs every 5 minutes. Fluorescence was quantified as the percent change in fluorescence relative to a baseline from -1 to 0 h prior to treatment. To determine if 17β-E_2_ increased neuro-seeER magnitude compared to control, we compared all data points from each experiment across 1 h windows between -1 to 0 h, 1-2 h, 3-4 h, 5-6 h, and 15-16 h.

### Photometry: Estrous Recordings

To capture the dynamics of the neural response to estrogen fluctuations across a full estrous cycle, we recorded neuro-seeER fluorescence using the FP3002 in intact female mice for at least 6 days. Estrous stage was estimated based on the relative proportion of different vaginal epithelial cell types observed following daily vaginal lavage, as described above. To compare neuro-seeER fluorescence across the estrous cycle, we trimmed the full recording to a 4 day period that included all cycle stages and quantified green channel fluorescence as the percent change relative to the mean signal magnitude across the 4 days being analyzed. To determine if neuro-seeER fluorescence differed across estrous stages, for each recording, we quantified and compared mean green fluorescence across metestrus (0-24 h), diestrus (24-48 h), proestrus (48-60 h), and estrus (60-96 h). Neuro-seeER signal correlations were computed between matched signals (*i.e.*, simultaneously recorded VMHvl and MPO neuro-seeER) or mismatched signals (*i.e.*, VMHvl and MPO neuro-seeER recorded during different experiments).

### Statistical Analysis

Statistical analyses were performed using GraphPad Prism Software (Version 10), in R (Version 4.4), or MATLAB (Version 2024b, 2025a). Statistical significance was defined as p < 0.05. All tests were two-tailed. Model assumptions were evaluated using standard diagnostic procedures. Full model outputs and N’s are provided in Table S1.

*In vitro* timelapse data: data were analyzed using linear mixed-effects models with hormone treatment, receptor transfection, time, and their interactions as fixed effects, and biological replicate as a random intercept. Models were fit using *lme4* (1.1), and significance of fixed effects was assessed with Type III Wald χ² tests using *car* (3.1). When significant hormone × receptor or higher-order interactions were detected, post-hoc comparisons were performed within each receptor condition, holding receptor constant, to evaluate hormone effects in receptor-positive versus control cells. Summary metrics derived from timelapse traces (AUC, time-to-maximum fluorescence, and initial rate of change) were analyzed with analogous mixed-effects models including hormone, receptor, and hormone × receptor as fixed effects and cell identity as a random intercept. Fixed-effect significance was evaluated with Type III Wald χ² tests, with post-hoc contrasts performed within receptor conditions when interactions were present.

Fixed *in vivo* fluorescence data: Quantitative induction of estrogen responsive neurons from ovariectomy and estrous-cycle experiments were analyzed using Firth-corrected logistic regression (brglm2) with treatment group or estrous stage as the fixed effect. Predefined contrasts (e.g., OVX vs. vehicle; each estrous stage vs. metestrus) were used to test specific hypotheses.

Steroid hormone levels: Group differences were assessed using Kruskal–Wallis test. Dunn’s comparisons were performed post-hoc.

Uterine weights: Group differences were assessed using Welch’s ANOVA. Brown–Forsythe tests were used to evaluate homogeneity of variance. Because variances were unequal, post-hoc pairwise comparisons were performed using Dunnett’s T3 multiple-comparisons test.

Live *in vivo* photometry experiments: To determine if 17β-E_2_ increased neuro-seeER signal magnitude relative to vehicle control, we used a linear mixed effects model with the formula ’DF/F ∼ Treatment * TimeWindow + (1|Experiment)’ to better handle its nested, longitudinal design. To compare neuro-seeER fluorescence across estrous stages, we used a repeated-measures ANOVA followed by multiple comparison correction with the Tukey-Kramer test. To compare latency of peak neuro-seeER signal, data were confirmed to be normal using a Shapiro-Wilk test and were compared using a paired t-test. To compare distributions of correlation coefficients, we used a 2-way Kolmogorov-Smirnov test. Data are presented as individual values with mean ± standard error of the mean (SEM).

## Data availability

Cell-by-cell quantification outputs as well as all images used for quantification of neuro-seeER fluorescence in figures 2 and 4 are available at DOI: 10.5061/dryad.xksn02vw4 All custom R packages used for data quantification and visualization are available at github.com/jevanveen

Raw and processed sequencing files are available at NCBI GEO under accession number GSE-312387

## Acknowledgements

Allisa Le (Princeton) for mouse husbandry and genotyping. Megan Massa and Alexa Wheeler for pioneering preliminary proof of concept studies with the neuro-seeER. The Broad Stem Cell Research Center (UCLA) for flow cytometry analyses and Incucyte™ experiments.

## Author Contributions

ALC conducted experiments, analyzed data, prepared figures, and wrote the manuscript. EMG acquired funding, designed and conducted experiments, analyzed data, prepared figures, and edited the manuscript. AM designed and performed in vitro experiments. ALH conducted LC-MS/MS experiments, analyzed data, and edited the manuscript. SY and IG conducted experiments and analyzed data. NPS and KA conducted experiments. KKS supervised LC-MS/MS studies and edited the manuscript. JEV designed seeER construct and performed quantification and bioinformatics analyses. JEV, ALF, and SMC acquired funding, designed and supervised the reporter expression analyses, and edited manuscript and figures.

---

**Fig. S1.**
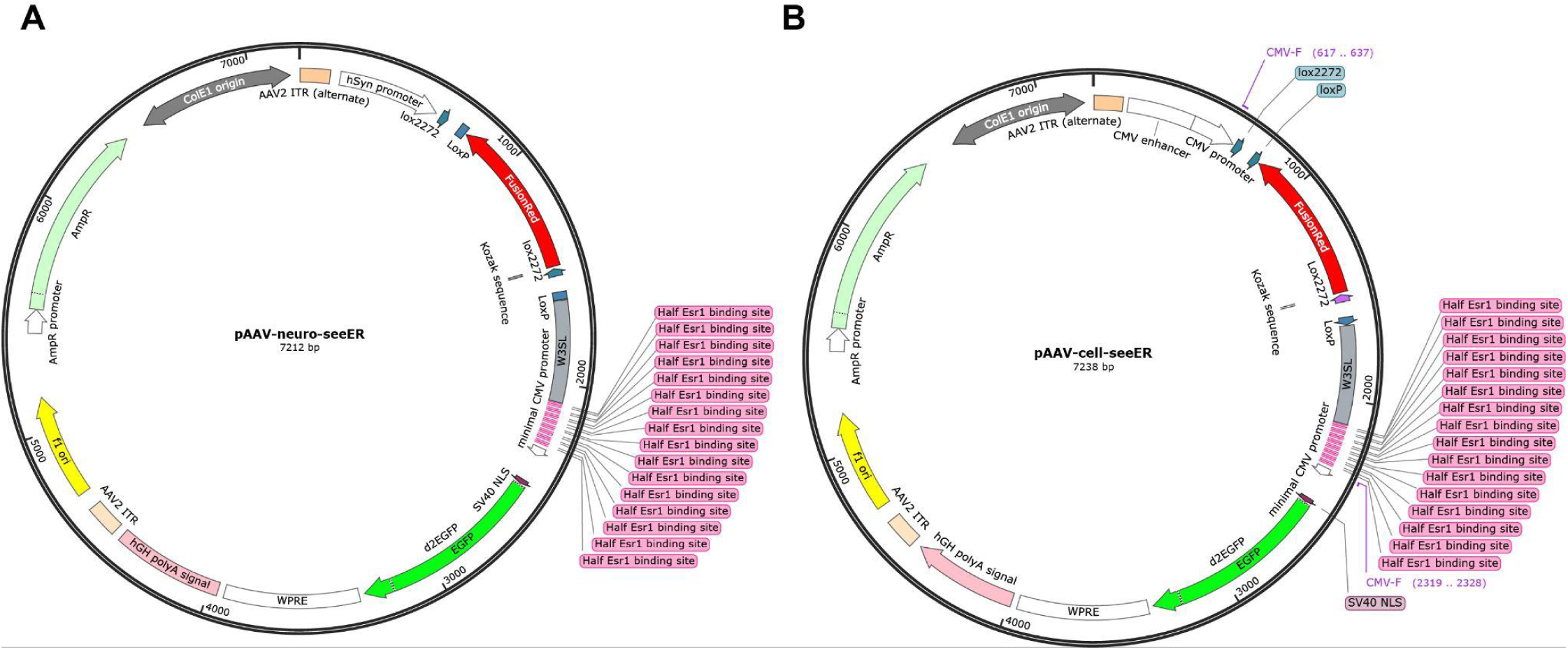
A, pAAV-neuro-seeER packaged in AAV virus and transduced into the hypothalami of Esr1-Cre mice. B, pAAV-cell-seeER transfected directly into 293T cells.

**Fig. S2.**
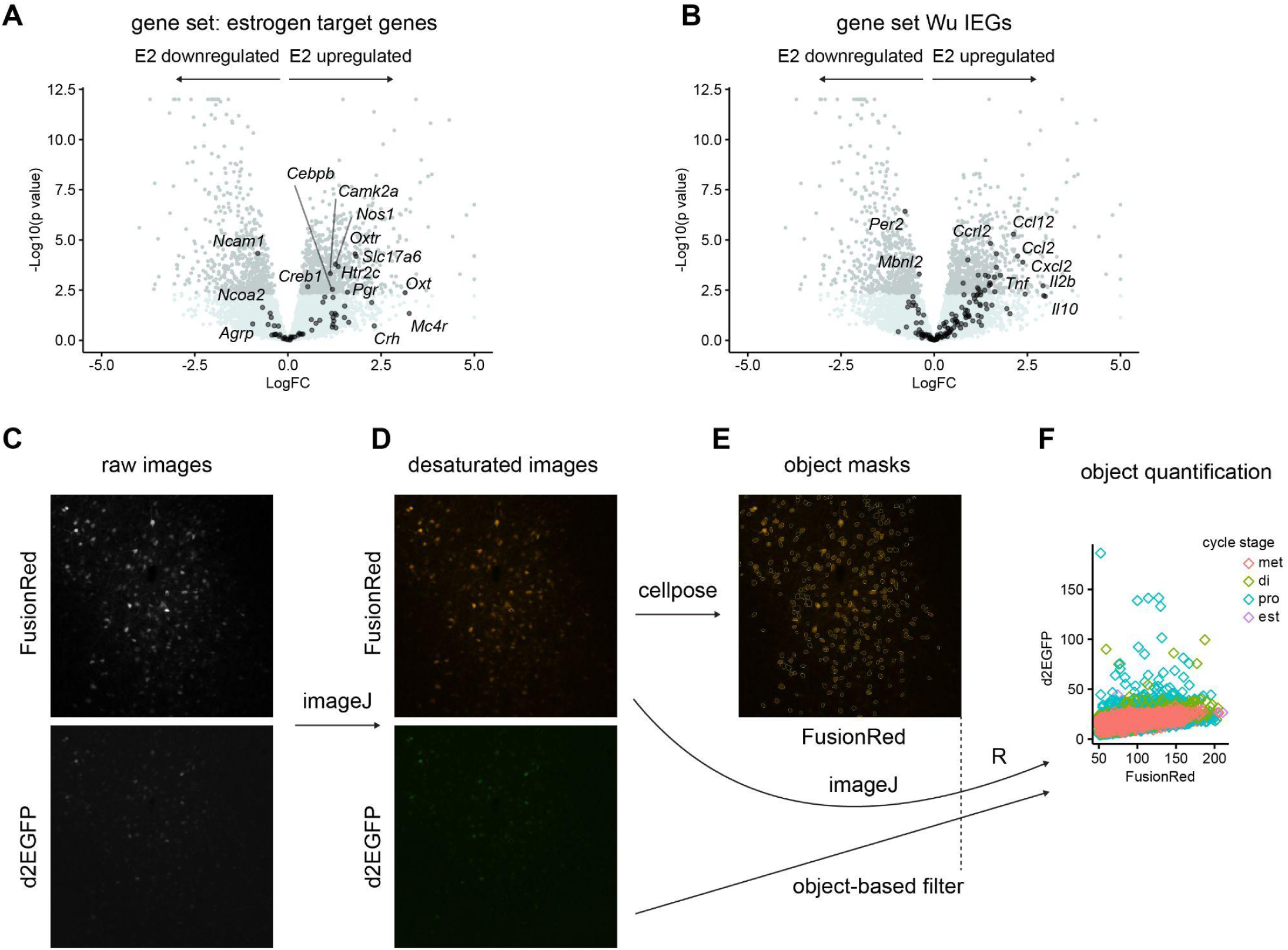
A, volcano plot of all genes differentially expressed between high-EGFP and low-EGFP cells in flow-seq analyses. Overlaid are genes reported in the literature to be transcriptional targets of E2 in the brain. B, volcano plot of all genes differentially expressed between high-EGFP and low-EGFP cells in flow-seq analyses. Overlaid are immediate early genes from^29^. C-F, image processing and analysis pipeline to identify estrogen responsive neurons. C, raw images are exported as tiffs. D, saturated pixels from either FusionRed or EGFP channels are removed from both images. E, neuronal objects are identified using cellpose-SAM^68^ on the FusionRed channel. F, unscaled, desaturated images are quantified at the individual object level and fluorescence is plotted and analyzed in R.

**Fig. S3.**
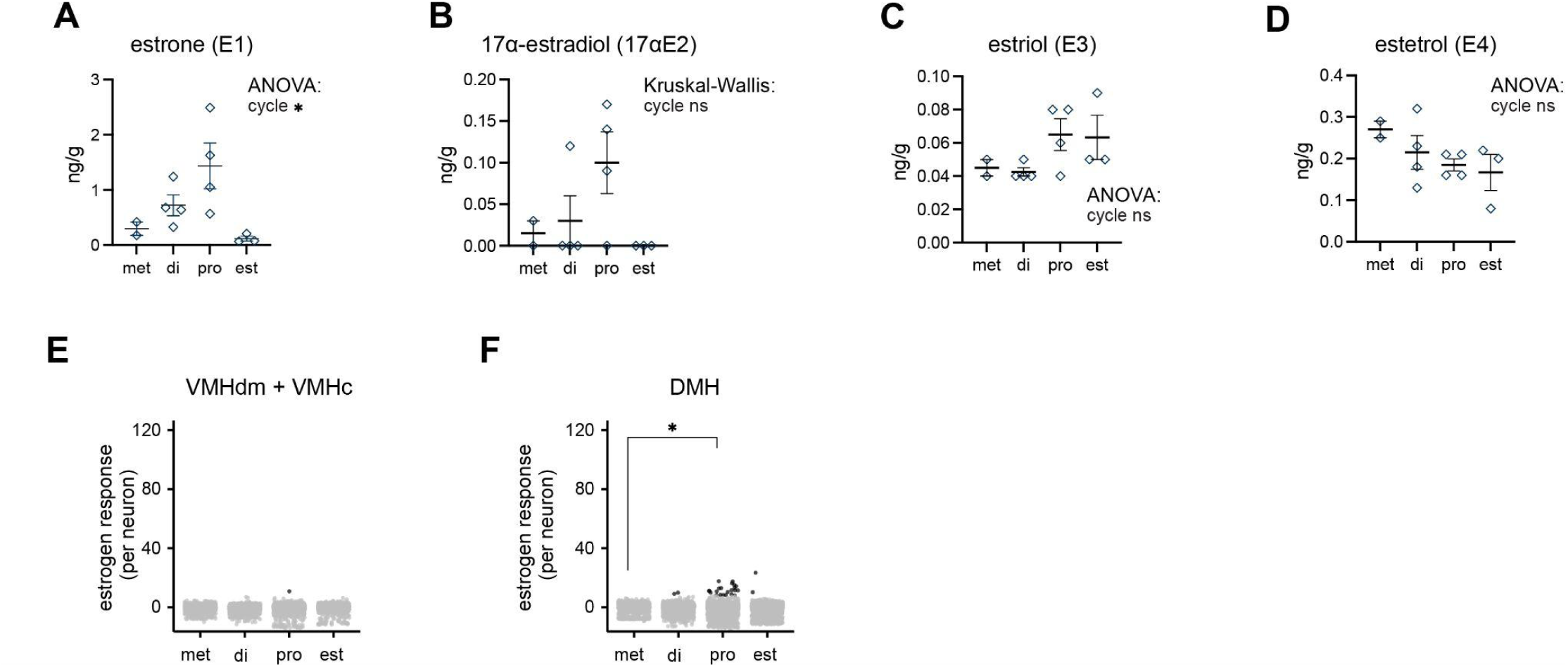
A, Ovarian estrone (E1, A), 17α-estradiol (17α-E2, B), estriol (E2, C), and estetrol (E4, D) levels across estrous cycle stages. E, Total number of estrogen responsive neurons (black) and averages per mouse (green) in the dorsomedial (VMHdm) + central (VMHc) portions of the VMH. F, Total number of estrogen responsive neurons (black) and averages per mouse (green) in the dorsomedial hypothalamus. Abbreviations: DMH, dorsomedial hypothalamus. VMH, ventromedial hypothalamus. met: metestrus, di: diestrus, pro: proestrus, est: estrus. *, p<0.05.

